# 3D Adaptive Optical Nanoscopy for Thick Specimen Imaging at sub-50 nm Resolution

**DOI:** 10.1101/2020.11.25.398958

**Authors:** Xiang Hao, Edward S. Allgeyer, Jacopo Antonello, Katherine Watters, Julianne A. Gerdes, Lena K. Schroeder, Francesca Bottanelli, Jiaxi Zhao, Phylicia Kidd, Mark D. Lessard, James E. Rothman, Lynn Cooley, Thomas Biederer, Martin J. Booth, Joerg Bewersdorf

## Abstract

Understanding cellular organization demands the best possible spatial resolution in all three dimensions (3D). In fluorescence microscopy, this is achieved by 4Pi nanoscopy methods that combine the concepts of using two opposing objectives for optimal diffraction-limited 3D resolution with switching fluorescent molecules between bright and dark states to break the diffraction limit. However, optical aberrations have limited these nanoscopes to thin samples and prevented their application in thick specimens. Here, we have developed a nanoscope that, by utilizing an advanced adaptive optics strategy, achieves sub-50 nm isotropic resolution of structures such as neuronal synapses and ring canals previously inaccessible in tissue.

---

The past two decades have witnessed a revolution in far-field light microscopy. The newly developed nanoscopes (or super-resolution microscopes)^1, 2^ are exceptional in their ability to noninvasively visualize cellular structures below the diffraction limit in all three dimensions (3D). This has enabled investigations of fundamental biological questions at the nanoscale with light microscopy for the first time. However, while nanoscopy can achieve 20 to 40 nm resolution laterally (*xy*), the axial (*z*) resolution in the axial dimension is at least two-times worse because of the limited illumination and/or detection aperture of single-objective systems.

To address this challenge, the axial resolution can be improved by coherently combining light from two opposing objectives with a common focus in a so-called ‘4Pi’ geometry^3, 4^. Interference in the common focal region dramatically sharpens the point-spread function (PSF) axially, improving the resolution 3- to 7-fold^5^. Further, collecting light with two objectives doubles the fluorescence detection efficiency. In combination with nanoscopy methods, such as stimulated emission depletion (STED) nanoscopy^6, 7^, reversible saturable optical linear fluorescence transitions (RESOLFT) nanoscopy^8^, or single-molecule switching (SMS; including (F)PALM, (d)STORM, GSDIM) nanoscopy^9-11^, 4Pi microscopes have realized nearly isotropic resolution of 20-50 nm in 3D. However, 4Pi microscopy is limited by axially repetitive side-lobes in the effective PSF, which can lead to ‘ghost images’ of structures above and below the focal plane. Further, the resolution is quickly compromised by accumulated optical aberrations when imaging deeper than a few microns off the coverslip. Because of these limitations, 4Pi nanoscopy applications were initially restricted to samples of <1 μm in thickness, and only recently have 4Pi nanoscopes been reported for whole-cell imaging^8, 11, 12^. However, 4Pi nanoscopy applications in tissues remain unexplored.

Here we present the development of a new nanoscope with sub-50 nm 3D resolution imaging capabilities to whole cells and tissue. A simplified schematic of our instrument, a laser-scanning STED nanoscope^13^ utilizing two opposing objective lenses which expands on a previously published isoSTED instrument by Schmidt *et al*.^6^, is shown in **Fig. 1a** (see **Supplementary Fig. S1** for the details). Fluorescence is restricted in all three dimensions to a sub-diffraction-sized volume by depleting excited fluorophores in the excitation volume’s periphery using stimulated emission. Switching off fluorophores in the periphery is performed by a depletion laser, which forms a hollow, sphere-shaped focus with a red-shifted wavelength, superimposed on the excitation focus. To generate a depletion profile for isotropic compression of the fluorescence emission volume, two depletion patterns featuring a common focal zero are combined: one for lateral (STED_*xy*_) and one for axial (STED_*z*_) depletion (**Fig. 1a, b**). While the STED_*xy*_ pattern is generated by the constructive interference of two ring-shaped foci, the STED_*z*_ beams interfere destructively, which in combination results in a “zero”-intensity minimum at the center surrounded by a high-intensity crest in all directions.

**Fig. 1.**
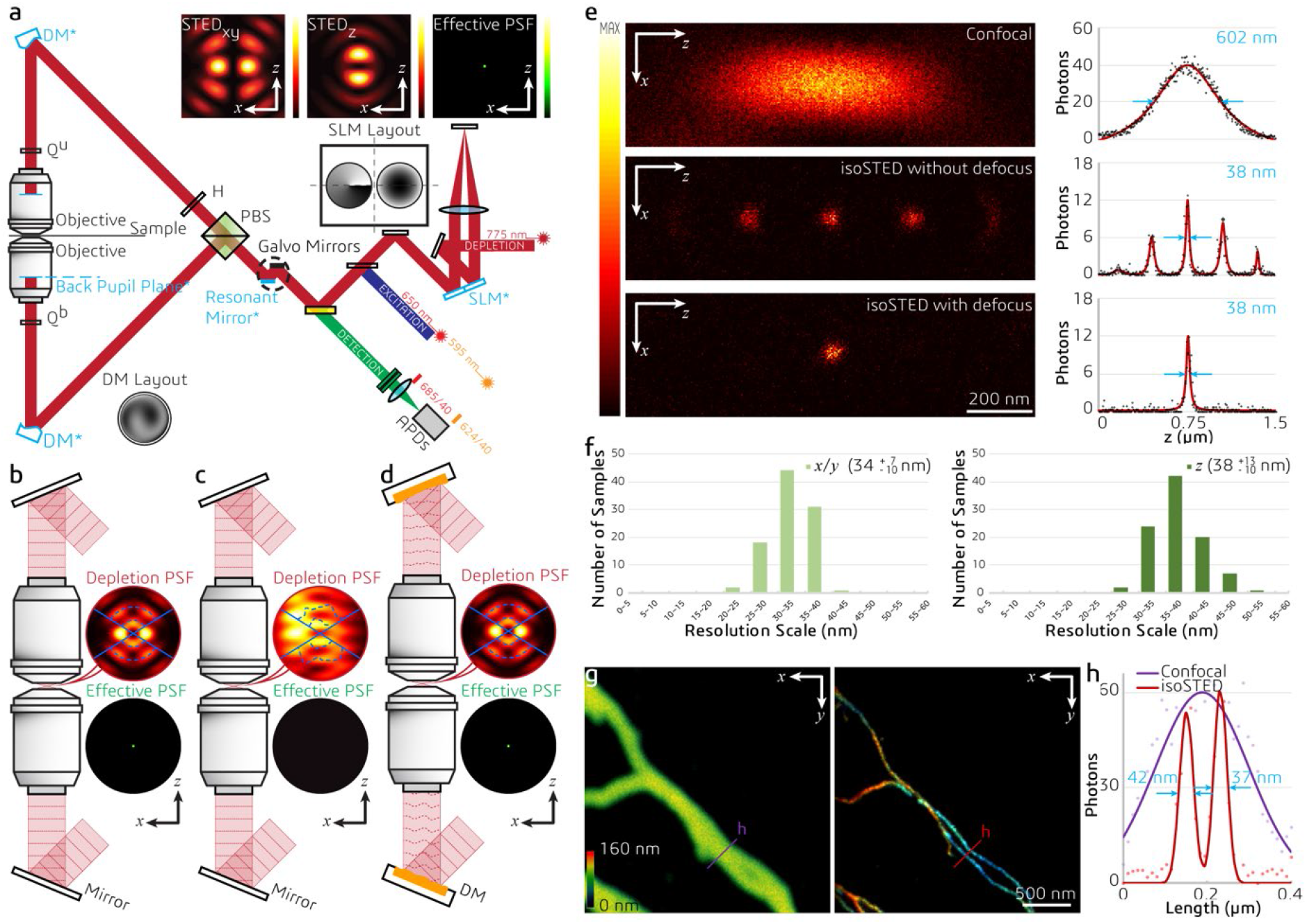
Principle and performance of adaptive optics (AO) isoSTED nanoscopy. (**a**) Simplified schematic. Abbreviations: PBS, polarizing beam splitter; H, half-wave plate; Q^u^, upper quarter-wave plate; Q^b^, bottom quarter-wave plate; DM, deformable mirror; SLM, spatial light modulator. The conjugated planes are indicated with an asterisk and highlighted in blue. Inserts on top: simulated intensity distributions in the focal region. (**b**) – (**d**) Principle of aberration correction in isoSTED nanoscope. Depletion and effective PSFs with and without sample-induced aberrations are shown as (**b**) and (**c**), respectively. (**d**) The deformable mirrors in the interference cavity introduce phase patterns that counteract aberrations, restoring nearly perfect optical performance. (**e**) *xz*|_*y*=0_ cross sections of the effective PSFs as measured with a 20-nm diameter crimson fluorescent bead. From top to bottom: single-objective confocal PSF, isoSTED PSFs with (bottom) and without (middle) defocus added to the STED_*z*_ pattern. To the right of each PSF, the respective axial intensity profiles (*z*) are displayed. The raw data (black dots) are fitted with a Lorentzian function (red curves), and the corresponding FWHMs are highlighted in blue. (**f**) Histogram of the lateral (left) and axial (right) FWHMs in isoSTED mode. At the top-right corner of each histogram, the average FWHM and the corresponding value range are given. (**g**) Confocal and isoSTED images of microtubules in the same region of a COS-7 cell. The distribution of α-tubulin (labeled with ATTO 647N) is visualized. The axial position of the microtubules is indicated using a rainbow colormap. A total of 16 optical sections (256×256 pixels each) were imaged. To maintain comparable brightness and contrast, the pixel dwell time in the confocal image (left) is 6 times shorter than that in isoSTED image (right). (**h**) Intensity profiles at the positions denoted by the solid lines in (**g**). The raw data (dots) are fitted with Lorentzian functions (solid curve). The corresponding FWHMs of the red line profiles from the isoSTED image are highlighted in blue.

To enable sub-50-nm isotropic resolution in thick specimens, our isoSTED nanoscope is built around two 100×, 1.35NA, silicone-oil objective lenses in a vertically oriented 4Pi interference cavity (see **Methods** and **Supplementary Fig. S1**). For multi-color imaging, two excitation beams and one depletion beam were supplied at 594, 650, and 775 nm wavelength, respectively, with an 80-MHz repetition rate. A 16-kHz resonant scanning mirror and two synchronized galvanometer mirrors were combined to enable fast 2D beam scanning.

Our new design provides three major improvements over the original isoSTED nanoscope^6^. First, using orthogonal polarization components from the same laser to generate the STED_*xy*_ and STED_*z*_ depletion patterns (**Supplementary Fig. S1**) leads to intrinsic co-alignment of the two depletion beams. To prevent these patterns from interfering with each other, we introduced a pulse delay module whose layout is inspired by a Michelson interferometer (**Supplementary Fig. S1**). A double-pass spatial light modulator (SLM) configuration^14^ was used to encode separate phase profiles upon each polarization component. Second, we added two quarter-wave plates in front of the objective back apertures to convert the linear polarization of the laser sources to circular polarization. This minimized excitation and depletion selectivity with respect to fluorophore dipole orientation and also reduced laser-induced background in the recorded images (see **Methods, Supplementary Fig. S2** and **S3**). Third, two deformable mirrors (DMs) were respectively placed into the upper and lower beam paths (**Fig. 1a-d**). Aberrations were corrected using a sensorless adaptive optics (AO) architecture^15^. A previously developed 4Pi aberration model^16^ enabled us to consider the whole isoSTED nanoscope as a single optical system and simultaneously optimize the wavefronts for both objectives using a single set of coefficients to achieve an aberration-free focus. Additionally, these DMs allow a small amount of defocus to be applied in both objective beam paths. This additional defocus virtually eliminates the repetitive axial side-lobes inherent to 4Pi microscopy^17^.

To quantify the resolution of our system, we imaged ninety-seven 20-nm diameter far-red fluorescent beads (**Fig. 1e**). In confocal mode, the full-width-at-half-maximum (FWHM) of the PSF was measured as 216 and 606 nm in the lateral and axial directions, respectively. Switching on the STED depletion beam resulted in an isotropic effective PSF of ∼35 nm (average over all the beads, **Fig. 1f** and **Supplementary Fig. S4**). In good agreement with simulations^17^, when no defocus was added to the STED_*z*_ beam, fluorescence was detected from the primary (>50% of the main peak) and secondary (10-20% of the main peak) side-lobes of the effective PSF. Next, we tested the resolution in COS-7 cells with ATTO 647N-immunolabeled microtubules (**Fig. 1g**). Example intensity profiles from imaged microtubules are presented in **Fig. 1h**. The system resolution was quantified from 56 intensity profiles by nested-loop ensemble PSF fitting^18^ at ∼39 nm.

Optical aberrations are a critical issue affecting 4Pi microscopy. Under ideal conditions, the effective PSF of our system is virtually side-lobe free after adding moderate defocus to the STED_*z*_ pattern (**Fig. 1e**) as mentioned above. In practice, however, the refractive index (RI) mismatch between the silicone oil and the mounting medium results in depth-dependent aberrations that subsequently result in depth-dependent side-lobes (**Supplementary Fig. S5**). These side-lobes manifest themselves as ‘ghost images’ above and below the real features in an image. In addition, when the sample is scanned in the *z*-direction, the positions of the PSF interference maxima shift under the PSF envelope, rendering any deconvolution procedure based on a spatially invariant PSF essentially useless. In principle, deconvolution with a variable PSF^19^ can mitigate this phenomenon, but still relies on an accurate knowledge of the PSF, and is limited to images with good signal-to-noise ratio (SNR). Therefore, a more effective approach is to minimize sample-induced aberrations using AO. Using the 4Pi aberration model and sensorless AO strategy mentioned above (see also **Methods**), sample-induced aberrations were corrected and a virtually side-lobe free PSF was restored. This strategy enabled us to image fine subcellular structures in one and two colors (**Supplementary Fig. S6**) within the cells, for example, microtubules in COS-7 cells (**Fig. 1g**), synaptonemal complexes in mouse spermatocytes (**Fig. 2a – d** and **Supplementary Video S1**), the Golgi apparatus in HeLa cells (**Fig. 2e–i** and **Supplementary Video S2**), and the endoplasmic reticulum (ER) and mitochondria in COS-7 cells (**Fig. 2j–l** and **Supplementary Video S3**).

**Fig. 2.**
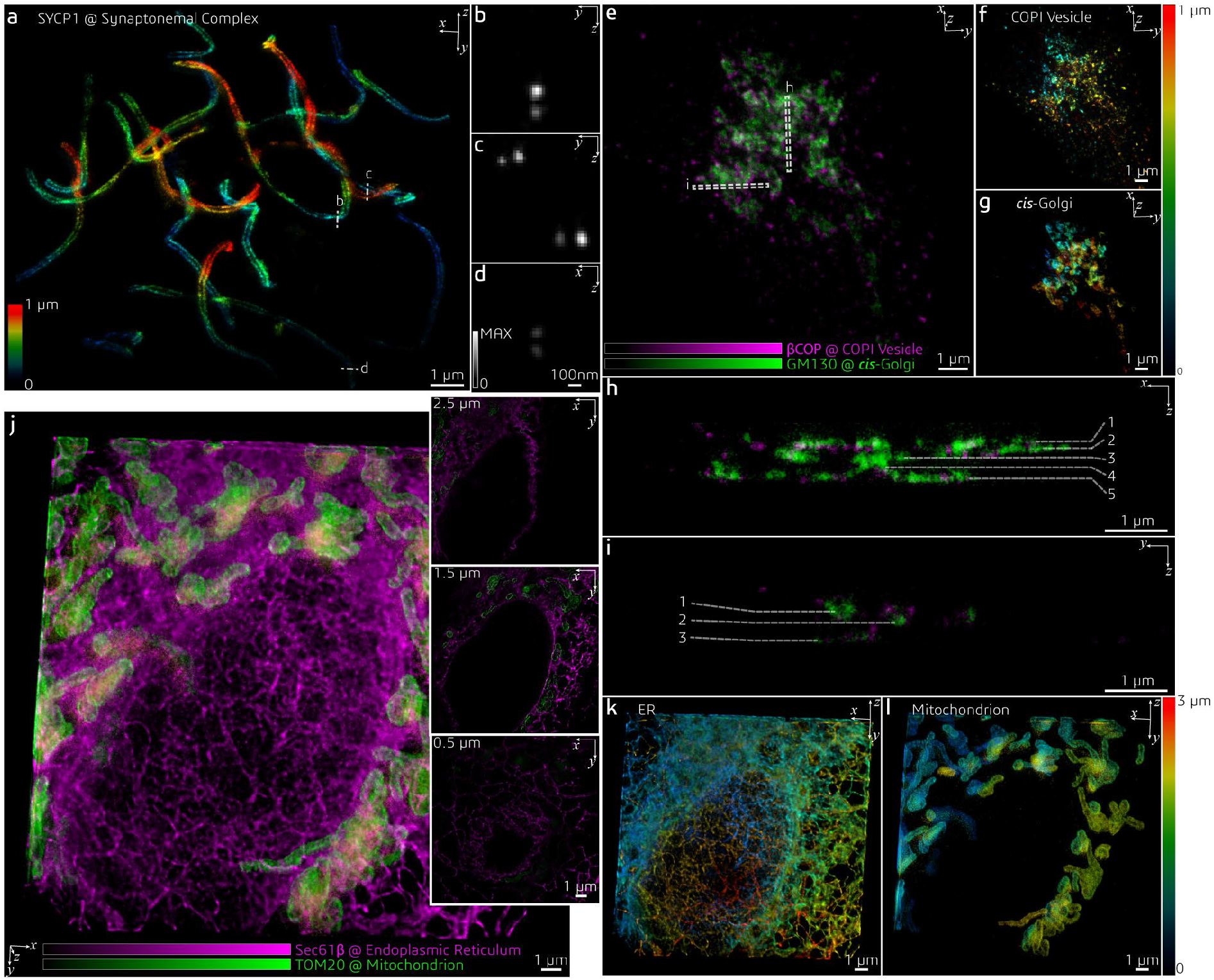
Applications of AO isoSTED nanoscopy in cultured mammalian cells. (**a**) Synaptonemal complexes in a mouse spermatocyte visualized by immunolabeling SYCP1 with ATTO 647N. (**b**) – (**d**) Cross sections of 1×1 μm^2^ regions at the positions denoted by the dashed line in (**a**). The images are normalized to the same peak intensity. (**b**) – (**d**) were smoothed using a 2D Gaussian kernel with a standard deviation of 1 pixel. (**e**) COPI vesicles (magenta) and Golgi apparatus (green) in a HeLa cell visualized by immunolabeling GM130 with ATTO 594 and βCOP with ATTO 647N. (**f**) - (**g**) Axial positions of vesicles (**f**) and *cis*-Golgi cisternae (**g**) in (**e**) indicated using a rainbow colormap. (**h**) - (**i**) Cross sections at the positions denoted by the dashed boxes in (**e**). Cisternae in the Golgi apparatus are indicated by the dashed lines. (**j**) ER (magenta) and mitochondria (green) in a COS-7 cell visualized by immunolabeling Sec61β with ATTO 594 and TOM20 with ATTO 647N. *xy* cross sections at different depths (*z*) are shown on the right. (**k**) - (**l**) Axial positions of mitochondria and ER in (**j**) indicated using a rainbow colormap.

Compared to typical mammalian cells cultured on coverslips (3-5 μm thick), RI inhomogeneities in tissue samples that are usually much thicker pose a greater challenge for aberration-sensitive techniques such as isoSTED nanoscopy. When mounting a sample for 4Pi imaging, a coverslip sandwich must be created with sufficient space to accommodate thick samples between the coverslips. While the distance between coverslips can be as low as 5 to 10 μm for samples of cultured cells, this distance must be expanded to >30 μm for most tissues to avoid compression artifacts (**Supplementary Fig. S7**). This increased sample thickness amplifies any existing RI mismatch between mounting medium and objective immersion liquid. To overcome this challenge and enable isoSTED nanoscopy in tissues, we developed a multistep method to optimize the DM surface shapes (see **Fig. 3a, Supplementary Fig. S8**, and **Methods**). First, after system aberrations were corrected, the two interference arms of the isoSTED instrument were treated as two separate microscopes^20, 21^, and the wavefront distortions in the upper and lower pupils were corrected separately by imaging gold beads attached to one of the coverslips in confocal. After this coarse aberration correction step, the focal plane was translated to the middle of the tissue. Next, we applied sensorless AO to minimize aberrations based on optimization of the fluorescence signal from the labeled biological structures. This correction step employed the 4Pi-aberration model previously developed^16^. The STED*xy* pattern was turned on, and a weak depletion power (∼10% of the power used for imaging) was chosen in order to minimize bleaching.

**Fig. 3.**
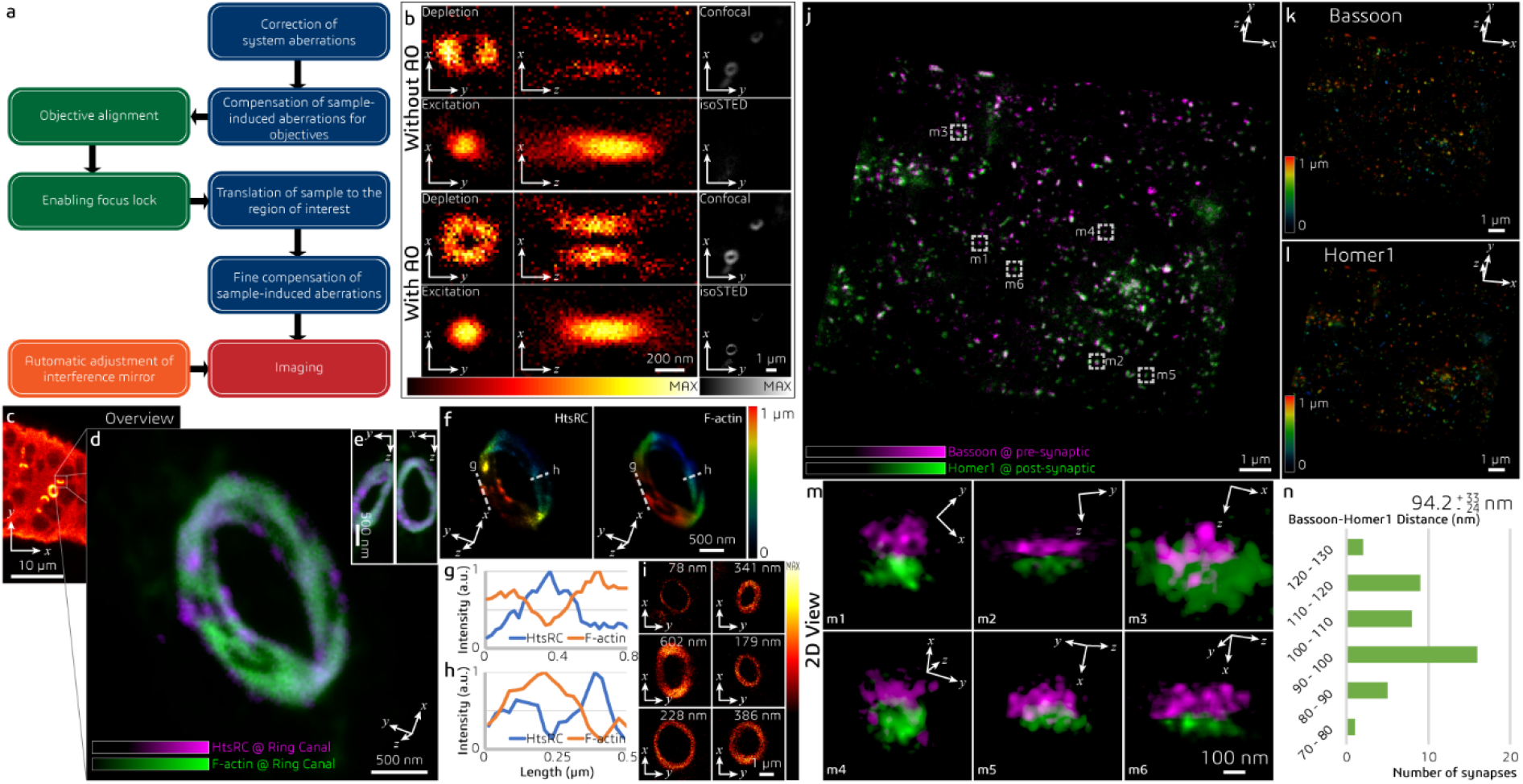
IsoSTED images of Drosophila egg chambers and mouse brain tissue. (**a**) Control loop of the AO strategy used for imaging tissue samples via isoSTED nanoscopy. (**b**) Effects of AO on the imaging performance of isoSTED. The figures in the left and middle columns show the *xy*|_*z* = 0_ (left) and *xz*|_y = 0_ (middle) cross-sections of the PSFs with (bottom two rows) and without (top two rows) AO aberration correction. The PSFs were obtained by imaging 150-nm-diameter gold beads. The right column shows 2D images of ring canals in a Drosophila egg chamber recorded in confocal and isoSTED modes. The F-actin proteins on the ring canals are immunolabeled with ATTO 594. (**c**) Overview image of Drosophila egg chambers. (**d**) Magnified image of a single ring canal recorded in isoSTED mode showing F-actin (green) and HtsRC (magenta) immunolabeled by ATTO 594 and ATTO 647N, respectively. (**e**) Front (left) and side (right) views of the ring canal shown in (**d**). (**f**) Axial positions of HtsRC (left) and F-actin (right) indicated using a rainbow colormap. (**g**) - (**h**) Intensity line profiles at the positions denoted by the dashed lines in (**f**). (**i**) F-actin staining of six ring canals in the same egg chamber. At the top-right corner of each sub-figure, the ring thickness is given (measured as the FWHM of the intensity profile in radial direction in the corresponding 2D ring). (**j**) IsoSTED image of hippocampal excitatory synapses visualized by immunolabeling the presynaptic active zone marker Bassoon and the postsynaptic scaffolding molecule Homer1 with ATTO 594 and ATTO 647N, respectively. (**k**) - (**l**), Axial positions of Bassoon (**k**) and Homer1 (**l**) in (**j**) indicated using a rainbow colormap. (**m**) Magnified projection images of the pre- and postsynaptic labels at the positions highlighted in (**j**). The trans-synaptic axes in these figures are rotated into the viewing planes. (**n**) Histogram of the Bassoon-Homer1 distances (measured as the peak-to-peak distance between the intensity distributions of the two stainings). At the top-right corner of the histogram, the average distance and the value range are given.

We demonstrate this multistep AO correction on ring canals, the cytoplasmic bridges that form from cells with arrested mitotic cleavage furrows in a Drosophila egg chamber (**Fig. 3c** and **Supplementary Video S4**). The proteins HtsRC and F-actin, both localized at the ring canals, were immunolabeled with ATTO 647N and ATTO 594, respectively. We compared the effects of AO in the same field of view in both confocal and isoSTED modes (**Fig. 3b**). While our AO approach somewhat improved the image quality in confocal, the improvements in isoSTED mode were striking: applying our aberration correction in isoSTED mode increased the image quality from essentially useless (**Fig. 3b**, second row) to a level where nanoscale features of the ring canals became clearly visible (**Fig. 3d-i, Fig. 3b**, forth row). The 2-color 3D data set of a ∼1.2-μm-diameter ring canal clearly reveals the different distributions of HtsRC and F-actin stainings (**Fig. 3d-f**) showing a more sparse, localized distribution of the HtsRC and a more continuous one for the F-actin. At the high 3D resolution of isoSTED nanoscopy, the two staining patterns show only little overlap (**Fig. 3f-h**). IsoSTED images of six different ring canals of similar diameter in the same egg chamber reveal notably different ring thicknesses in their HtsRC staining (**Fig. 3i**), ranging from sub-100 nm to ∼600 nm.

To demonstrate our approach on another challenging target, we imaged excitatory synapses in a 30-μm-thick mouse brain slice (**Fig. 3j** and **Supplementary Video S5**). A pair of pre- and postsynaptic proteins, the active zone marker Bassoon and the scaffolding protein Homer1 that labels excitatory synapses, were immuno-stained and detected with ATTO 594 and ATTO 647N-conjugated secondary antibodies, respectively. IsoSTED images (**Fig. 3k-m**) resolved the two protein distributions as juxtaposed, clearly resolved pairs, demonstrating the quantification capabilities of our instrument at the sub-diffraction-limit scale inside tissue. The mean distance between the pre- and post-synaptic markers was ∼100 nm, (**Fig. 3n**), a distance in excellent agreement with a previous report^22^ but now reported from much thicker sections.

In summary, AO isoSTED nanoscopy allows us to image fine structures with sub-50-nm isotropic resolution and demonstrates AO for nanoscopy techniques in thick samples. Stimulated emission is only one of several mechanisms to realize reversible fluorescence inhibition. Thus, the scheme presented in this work is generally applicable to any AO microscope with a 4Pi cavity. This underscores the potential of applying our AO strategy to other nanoscopy techniques, such as 4Pi-RESOLFT^8^ and 4Pi-SMS^10, 11^. AO has many potential applications beyond “simple” aberration correction in 4Pi techniques - suppressing 4Pi sidelobes by introducing defocus modes to the depletion wavefronts is just one possibility. More advanced wavefront control can be applied to customize PSFs and generate, for example, multiple foci for multifocus scanning in live-cell applications.

Combining expansion microscopy^23^ with isoSTED nanoscopy is an exciting new frontier which could achieve sub-10-nanometer isotropic resolution – a value which approaches the size of the labels themselves. AO will be instrumental in correcting for the depth-dependent aberrations stemming from the refractive index mismatch between the immersion medium of the objective and the hydrogel.

In conclusion, our development provides access to the 3D organization of tissue at the nanoscale by fluorescence microscopy and represents an important step towards understanding sub-cellular organization in the context of tissue.

## Methods

### Nanoscope setup

Design details are provided as Supplementary Information (**Supplementary Fig. S1**). In brief, the 4Pi interference cavity of the isoSTED system is set up vertically around two 100×, 1.35NA, silicone-oil objective lenses (UPLSAPO 100XS, Olympus). An 80-MHz pulsed depletion laser with a pulse length of ∼600 ps (775 nm, Katana HP, OneFive) is coupled into a 10-m long, polarization-maintaining single-mode fiber (PM-S630-HP, Nufern). One polarization component of the laser is delayed by a Michelson interferometer with respect to the orthogonally polarized component to avoid later interference between the two polarization components. The beam emerging from the fiber is collimated and illuminates a spatial light modulator (SLM, X10468-02, Hamamatsu), which applies a vortex and a moderate defocus phase pattern to the two orthogonally oriented polarization components of the depletion beam, respectively^17^. The SLM is imaged onto a 16-kHz resonance scan mirror (SC-30, EOPC), which is, in turn, imaged into the back pupils of both objectives.

Pulsed excitation light from two laser lines at wavelengths of 650 nm (LDH-P-C-650m, PicoQuant) and 594 nm (LDH-D-TA-595, PicoQuant) is coupled into the same 2-m long polarization maintaining single-mode fiber (P1-488PM-FC-2, Thorlabs) and merged with the STED beam path via a dichroic mirror (T740lpxr, Chroma). The polarization orientation of the excitation beam is tuned by a quarter-wave plate (QWP, AQWP05M-600, Thorlabs), a linear polarizer (LPVISE100-A, Thorlabs), and a half-wave plate (HWP, AHWP10M-600, Thorlabs) to an angle of 45° with respect to that of the depletion beam.

The resonance mirror scans the combined laser beams along the fast scanning axis (*x*), while two synchronized galvanometer scanning mirrors (dynAXIS XS, SCANLAB) scan the beams along the slow scanning axis (*y*). Together, the two galvanometer mirrors behave as a single scan mirror at the conjugate pupil plane, allowing the three scanning mirrors to act as a fast, dual-axis scanning system positioned in a plane conjugate to the pupil planes of the objectives. A polarizing beam-splitter (PBS, PBSH-450-00-100, CVI Laser Optics) is used to produce the counter-propagating pair of beams in the interference cavity. A HWP (AHWP10M-600, Thorlabs) before the PBS rotates the polarization orientations of the incident depletion beam to ±45° with respect to the polarization axis of the PBS, enabling precise 50/50 power allocation between the two arms of the cavity. Because of its incident 45° linear polarization, the excitation beam is rotated to 0° by the HWP in front of the PBS and is therefore only coupled into the upper arm of the interference cavity. The polarization of the beams in the upper arm is further rotated by another HWP (AHWP10M-600, Thorlabs) for later interference in the combined focus (**Supplementary Fig. S2**). The phase difference between the two arms is minimized by translating the BK-7 glass wedge (custom design, OptoCity) in the lower arm. In both arms of the interference cavity, a deformable mirror (DM, MultiDM-5.5, Boston Micromachines) is added in the plane conjugated to the back pupil of the corresponding objective, which allows for aberration correction and manipulation of the PSF. Circular polarization of all beams in the focus is ensured by QWPs (AHWP05M-600, Thorlabs) in front of the objectives.

The sample is mounted at the common focal plane of the objectives. The fluorescence from the sample is collected by both objectives, combined at the PBS, de-scanned by the scanning mirrors, and then separated from the common beam path via a custom-made, 5-mm-thick quad-bandpass dichroic mirror (ZT485/595/640/775rpc, Chroma). An additional dichroic mirror (ZT640rdc, Chroma) splits the fluorescence into two detection bands for multicolor imaging. For each band, two filters of the same type (separated by about 50 mm and mounted at a small angle relative to each other to avoid interference between them) reject stray excitation and depletion laser light (FF02-685/40-25, FF01-624/40-25, Semrock). Fluorescence in each detection band is coupled into a multimode stainless tubing fiber (FG105LCA, Thorlabs) acting as a confocal pinhole, with a pinhole size of ∼0.8 Airy units. Each fiber is connected to a single-photon counting avalanche photodiode (APD, SPCM-ACRH-13-FC, Excelitas). The measured fluorescence signals from the APDs are relayed to circuit boards for gated detection (custom built, Opsero Electronics). These gate boards are synchronized to the trigger signal from the depletion laser. The gate electronics allow software control of the detection window length and position. The same trigger signal and circuit boards are used to trigger the excitation lasers at a user-defined delay. Finally, the APD signal is acquired by a field-programmable gate array board (PCIe-7852R, National Instruments), which is synchronized to the resonance mirror. The system is controlled via a custom-made interface programmed in LabVIEW 2014 (National Instruments). The system allows line-by-line image acquisition.

### Adaptive Optics Strategy

#### Characterization of DMs

Each DM is characterized offline^24^ using an external Michelson interferometer setup before it is inserted into the 4Pi cavity. By tilting the flat mirror in the reference arm of the interferometer setup, the phase induced by the DM can be obtained using Fourier fringe analysis^25^ and phase unwrapping^26^. The characterization of the DM consists of computing the influence matrix *H* that maps a vector *u* containing the voltage of each actuator of the DM to the corresponding vector *z* that contains the coefficients of the Zernike analysis of the phase. The matrix *H* is computed by collecting a set of input-output vector pairs *u* and *z*, and by solving a least-squares problem^27^. After the DMs are installed into the 4Pi cavity, they are operated in open-loop and controlled using the vector of Zernike coefficients *z* as the independent variable, i.e., given a vector of desired Zernike coefficients *z*, the voltage vector *u* to be applied to the DM is found by minimizing the norm squared of *z* −*H*.*u*.

#### Aberration Control in the 4Pi Cavity

We employ the modes 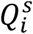 defined in Ref. 16 to model the aberration in the 4Pi cavity and to apply the aberration correction using a single set of coefficients 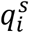. To operate the two DMs simultaneously, we relate these coefficients 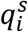to the vectors of Zernike coefficients *z*_*u*_ and *z*_*l*_, which correspond to the upper and lower pupil of the 4Pi objectives, respectively. The voltage to be applied to each actuator of a single DM is subsequently determined from either *z*_*u*_ or *z*_*l*_ as described in the previous section.

#### Aberration Correction

The flowchart used for aberration correction is presented in **Fig. 3a**. We employ an image-based, quadratic aberration-correction algorithm to apply the aberration correction at the three different steps outlined in the flowchart (See **Supplementary Fig. S8**). The algorithm involves taking a series of images where different amounts of a certain aberration mode are applied to the DMs while acquiring each image^15^. An image quality metric specified below is used to quantify the quality of each acquired image. By fitting a curve through the determined image quality metric values versus the amounts of the applied aberration, the optimal value that maximizes the image quality can be found. This process is repeated for each of the considered aberration modes.

The first step involves removing the system aberrations, i.e., the aberrations caused by misalignment of the optical components comprising the nanoscope and their manufacturing tolerances. In this step, we image 100-nm-diameter crimson fluorescent beads as the target. We optimize the setting of the objective correction collar, the SLM, and then we apply the image-based aberration correction to optimize the Zernike coefficients of the upper and lower pupils individually. The brightness of the image is taken for the metric.

In the second step, the two DMs are still independently adjusted to coarsely compensate sample-induced aberrations and to correct for the large amounts of aberrations caused by the refractive index (RI) mismatch between sample, embedding medium and immersion liquid (silicone oil). In this step, we image 150-nm-diameter gold beads^21^ attached to the bottom coverslip using the depletion beam for illumination. We optimize the first 11 Zernike coefficients in each pupil using the brightness and the PSF shape as the metric. To reduce the influence of noise, a threshold is set manually to remove background pixels. After this coarse aberration correction step, the objectives are aligned and locked (see “Focus-Lock Module” section).

In the final step, we optimize the 4Pi modes by simultaneously adjusting both DMs. In this step, we switch to 2D isoSTED mode and move to the area of interest within the tissue. At this step the fluorescence emission signal is recorded to apply the aberration correction. We optimize the following 4Pi modes 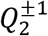 up to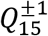. Note that in this step we do not consider the mode 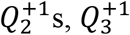, and 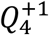, which cause displacement of the 4Pi PSF but do not affect the image quality^6^. In addition, mode 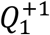 is corrected by translating a piezo-driven mirror in the 4Pi cavity (see **Supplementary Information Fig. S1**) instead of using the DMs. To ensure that the sensorless algorithm is sensitive to aberrations that affect the STED resolution, we activate the STED_*xy*_ phase pattern at a weak depletion power (∼10% of the power level used for imaging). This step is optional when imaging relatively thin cell samples, especially if the imaging volume is close to the coverslip. In contrast, for tissue samples, this step is essential to allow for high-quality isoSTED imaging. If necessary, the three stages are repeated to further improve the image quality.

To acquire a large imaging volume (>1 μm in *z*-direction), a linear bias compensation of mode 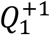(plus, optionally, defocus 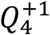 and 1^st^-order spherical aberration 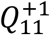) is applied when the sample is scanned in *z*-direction. The bias step size is manually selected to achieve constructive interference in the focal plane of the STED_*xy*_ focus throughout the whole imaging volume in the *z*-direction.

### Biological Sample Preparation

### Cell culture

COS-7 (ATCC, CRL-1651) and HeLa (ATCC, CCL-2) cells were grown in DMEM (Gibco, 21063029) supplemented with 10% fetal bovine serum (Gibco, 10438026) at 37 °C with 5% CO_2_.

### Microtubules

COS-7 cells were grown on #1.5, 25-mm-diameter round precision coverslips. Cells were rinsed 3 times with pre-warmed 1× PBS (37 °C) and then pre-extracted with 0.2% saponin in CBS buffer (10 mM MES, 138 mM KCl, 3 mM MgCl, 2 mM EGTA, 32 mM sucrose). After 1 min, the pre-extraction buffer was removed and the cells were fixed in pre-warmed (37 °C) 3% paraformaldehyde, 0.1% glutaraldehyde in CBS buffer for 15 min at room temperature. The cells were then washed with 1× PBS 3 times and immunofluorescence staining was performed using mouse anti-alpha tubulin primary antibody (T5168, Sigma Aldrich) at 1:1000 dilution, and goat anti-mouse ATTO 647N secondary antibody (50815, Sigma Aldrich) at 1:1000.

### Synaptonemal Complex

All experimental procedures involving the use of mice were performed in agreement with the Yale University Institutional Animal Care and Use Committee (IACUC). Testes (tunica removed) from 18-day old mouse pups were disrupted using forceps and a razor blade in 1 mL of 1× PBS with protease inhibitors (05896988001, Roche Complete Ultra). The cell suspension was then gently added to a 15-mL conical tube with 5 mL of 1× PBS with protease inhibitors and allowed to settle. After approximately 3 min, 5 1-mL aliquots of the cell suspension were places in 1.5-mL microcentrifuge tubes and centrifuged at 9000 RPM for 10 min. The supernatant was then aspirated, and the pellets were combined in 0.5 mL of 1× PBS per testes. 50-100 µL of the cell suspension were added to clean, #1.5, 25-mm-diameter round coverslips and allowed to sit for 30 min. The coverslips with cells were then fixed in 4% paraformaldehyde for 15 min at room temperature. The samples were then washed with 1× PBS 3 times, and immunofluorescence staining was performed using rabbit anti-SYCP3 primary antibody (ab15092, Abcam) at 1:1000 dilution, and goat anti-rabbit ATTO 647N secondary antibody (50815, Sigma Aldrich) at 1:1000.

### Golgi Apparatus

HeLa cells were fixed with 4% paraformaldehyde for 15 min at room temperature. Indirect immunofluorescence was carried out using anti-GM130 (BD Transduction Laboratories) and anti-ßCOP^28^ and secondary antibodies conjugated with ATTO 594 (76085, Sigma-Aldrich) and ATTO 647N (50815, Sigma Aldrich).

### ER + mitochondrion

COS-7 cells growing on #1.5, 25-mm-diameter round precision coverslip were transfected with mEmerald-Sec61-C-18, a gift from Michael Davidson (54249, Addgene plasmid) using Lipofectamine2000 (11668019, Invitrogen). The next day, the cells were fixed with fresh 3% paraformaldehyde (15710, Electron Microscopy Sciences), 0.1% glutaraldehyde (16019, Electron Microscopy Sciences) in PBS for 15 min at room temperature. The endoplasmic reticulum (ER) was labeled with mouse anti-GFP (A11120, Invitrogen), and mitochondria were labeled with rabbit anti-TOM20 (11415, Santa Cruz Biotechnology) primary antibodies at 4 °C overnight. Secondary antibodies anti-mouse ATTO 594 (76085, Sigma Aldrich) and anti-rabbit ATTO 647N (76085, Sigma Aldrich) were used at 1:1000.

#### Tissue Sample Preparation

### Drosophila egg chamber

Ovaries were dissected in IMADS buffer^29^ and fixed for 5 min in 4% paraformaldehyde in PBS with 0.6% Triton X-100. Fixed tissue was washed in PBT (phosphate-buffered saline with 0.6% Triton X-100 and 0.5% BSA) and incubated with 1:10 anti-HtsRC^30^. Secondary antibodies used were goat anti-mouse conjugated to ATTO 647N (Sigma-Aldrich, 1:500). F-actin was labeled with phalloidin conjugated to ATTO 594 (Sigma, 1:500).

### Mouse brain slice

All experiments were carried out in accordance with the recommendations of the NIH guidelines and the Tufts University Institutional Animal Care and Use Committee. The protocol was approved by the Tufts University Institutional Animal Care and Use Committee. Wild-type (C57BL6) adult male mice were transcardially perfused first with ice-cold PBS and then with 4% paraformaldehyde (in PBS, pH 7.4). Brains were isolated and post-fixed overnight in 4% PFA and subsequently washed and stored in PBS (all at 4°C). Brains were mounted in cold PBS and coronally sectioned at 30 µm using a vibrating microtome (Vibratome 1500, Harvard Apparatus). Sections were stored in PBS at 4 °C. Medial hippocampal slices were selected and washed four times for 15 min in PBS at room temperature. Next, non-specific antibody binding sites were blocked with 3% normal horse serum and 0.1% Triton-X 100 in PBS for 2 hours at room temperature. Tissues were incubated sequentially with primary and secondary antibodies at 4 °C for 48-72 hours and overnight, respectively. Presynaptic monoclonal mouse anti-Bassoon (SSP7F407, Enzo; 1:500) and postsynaptic polyclonal rabbit anti-Homer1 (160 003, Synaptic Systems GmbH; 1:500) were used as primaries. Goat anti-mouse IgG antibodies conjugated with ATTO 594 (76085, Sigma-Aldrich; 1:500) and goat anti-rabbit IgG antibodies labeled with ATTO 647N (40839, Sigma-Aldrich; 1:500) were used for secondaries. All primary and secondary antibodies were diluted in 3% normal horse serum and 0.1% Triton-X 100 in PBS. After the incubation steps, sections were extensively washed in PBS and then floated onto 25-mm-diameter precision glass coverslips in distilled water. All coverslips were first sonicated (Branson 2510 Ultrasonic Cleaner, Marshall Scientific) in 1 M potassium hydroxide for 15 min and then washed generously and allowed to dry. The coverslip opposite to the coverslip holding the specimen was treated with ∼80 μL of poly-L-lysine for 15 min at room temperature. After aspirating off the bead dilution, the coverslips were sandwiched into a mounting ring using mounting medium (CFM-3, Citifluor). A thin coat of clear nail polish was used to seal the coverslips into the mounting ring.

#### Sample Mount

All samples were mounted between two 25-mm-diameter, 170-μm-thick coverslips (CSHP-No1.5-25, Bioscience Tools) at 5-30 μm separation (see **Supplementary Fig. S7**). 150-nm-diameter gold beads were pre-deposited onto the bottom coverslip, acting as the reference for the objective alignment during the experiment. For fluorescent beads and cell samples, mounting medium (Prolong Diamond, Thermo Fisher Scientific) was applied to supply anti-fade protection and to narrow the RI difference between the sample and the silicone oil. After mounting, a custom-designed ring was used to hold the coverslips, and the gap between them was sealed using a two-component silicone (Twinsil 22, Picodent).

### Drift Correction

#### Focus-Lock Module

A focus-lock module is integrated into the nanoscope to stabilize the objective alignment. A 980-nm-wavelength CW laser is merged with the other beams via a dichroic mirror (ZT1064rdc-sp, Chroma) in the top arm of the interference cavity. The beam is focused by the upper objective, collected again by the lower objective and is separated from the common beam path by another dichroic mirror of the same type in the bottom arm. Before being focused by a 400-mm-focal-length lens onto a camera (DCC1545Mz, Thorlabs), the beam is modulated by a weak cylindrical lens (LJ1144RM-B, Thorlabs) to add astigmatism. The position and the shape of the focal spot are recorded, and monitored during the imaging. Specifically, the position of the spot reflects the relative positions of the objectives in *x* and *y* directions, while the spot shape is used for alignment in *z* direction. Spot position and shape are automatically analyzed and, when changes are detected, the piezo objective stages are translated to compensate for them using a proportional–integral–derivative (PID) control loop programmed in LabVIEW. The application of the focus-lock module enables long-term (>3 hours) imaging. Notably, as the laser power of the alignment laser is very low (< 5 μW), and the wavelength is far beyond the excitation spectrum, there is no visible impact to the sample or image when the focus-lock module is on.

#### Sample Drift

While sample drift is negligible in most experiments, cross-correlation is applied occasionally as an effective post-processing algorithm to correct for small sample drift. The whole dataset is split into *n* frames along the *z*-axis. The 2D cross-correlation is calculated between two contiguous frames, and the second frame is shifted accordingly to match the correlation peak and compensate sample drift in *x-* and *y*-directions.

### Image Processing and Visualization

Image data was recorded with custom-programmed LabVIEW software (see above). A 2D Gaussian smoothing filter (σ = 1 pixel) was applied to the data shown in Figure 2b. None of the other data was smoothed. No deconvolution algorithms were applied to any of the shown data. The freely available software package Fluorender (version 2.22.3, http://www.sci.utah.edu/software/fluorender.html) was used for 3D rendering.

## ACKNOWLEDGEMENTS

We thank Dr. George Sirinakis, now at the University of Cambridge, Dr. Yongdeng Zhang, Andrew E. S. Barentine from Yale University, Dr. Mary Grace Velasco, now at Abberior Instruments, and Dr. Emil B. Kromann, now at the Technical University of Denmark, for sharing the source code and fruitful discussions. We also thank Dr. Yong Wang at the University of Utah for technical support with FluoRender. This work was supported by the Wellcome Trust (095927/A/11/Z, 095927/B/11/Z, 203285/B/16/Z, 203285/C/16/Z), the G. Harold & Leila Y. Mathers Foundation, the National Institutes of Health (P30 DK045735 to Robert Sherwin; R01 GM043301 to L.C.) and the European Research Council (AdOMiS, N<u>o</u>. 695140).

## Author contributions

X. H., E. A., and J. B. designed the instrument. X. H., E. A., and J. A. built the instrument and developed the software. X. H., J. A., J. Z., and M. B. developed and optimized the adaptive optics strategy. X. H. and E. A. optimized and tested the instrument. K. W., J. G., L. S., F. B., P. K., M. L., J. R., L. C., T. B., and J. B. designed the biological experiments. K. W., J. G., L. S., F. B., P. K., and M. L. optimized the sample preparation protocols and prepared the samples. X. H., M. L., and J. B. visualized the data. All authors contributed to writing the manuscript.

## Additional information

Supplementary information is available in the online version of the paper. Reprints and permissions information is available online at www.nature.com/reprints. Correspondence and requests for materials should be addressed to J. B.

## Data Availability

The datasets generated during and/or analyzed during the current study are available from J. B. (the corresponding author) on reasonable request.

## Code Availability

The source code is available from J. B. (the corresponding author) on reasonable request.

