## Supplemental Information for "3D Adaptive Optical Nanoscopy for Thick Specimen Imaging at sub-50 nm Resolution"

### Supplementary Information

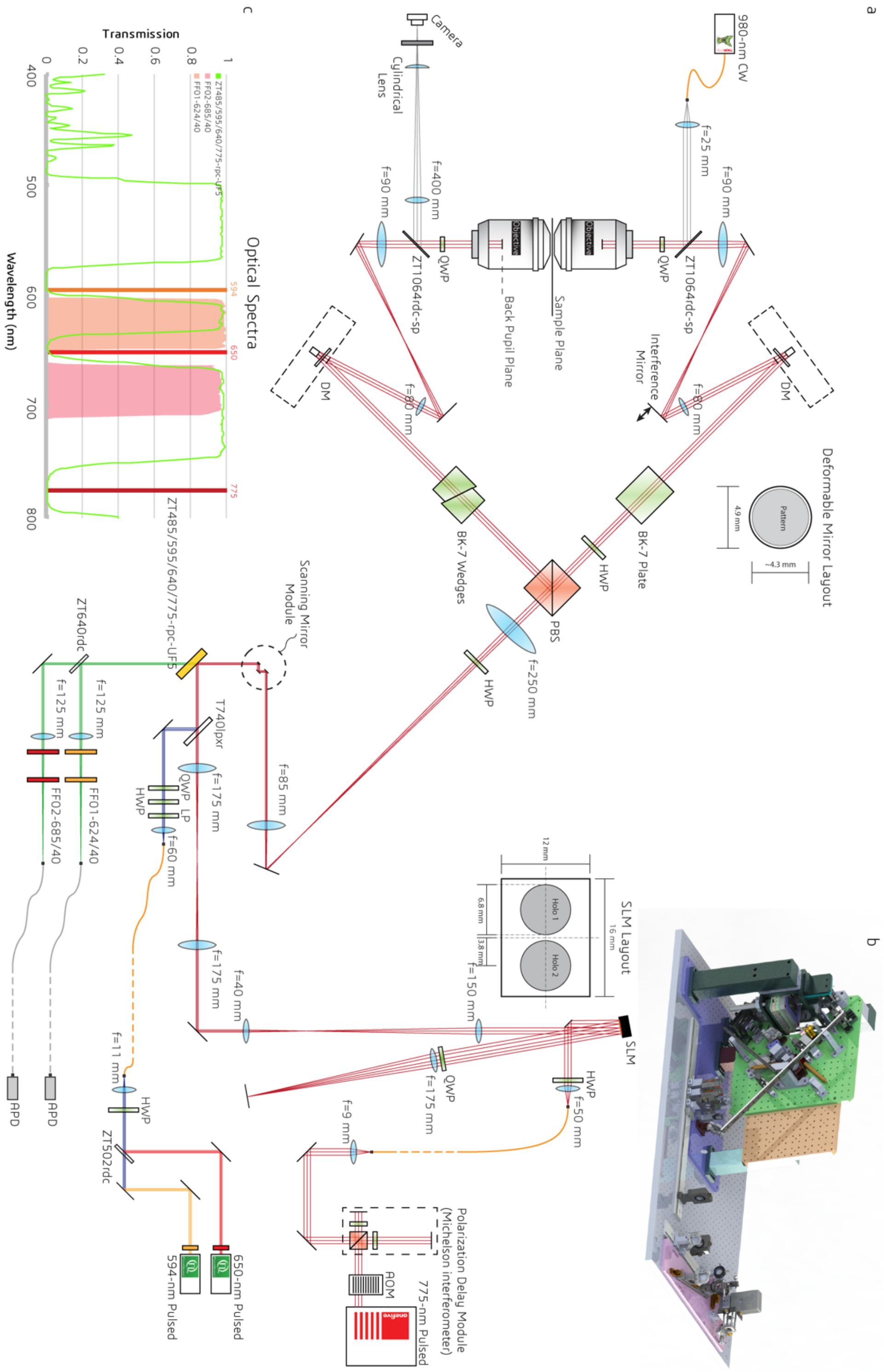


**Supplementary Fig. S1 | Design of isoSTED nanoscope.** (**a**) System layout. The system is built from the following components: excitation lasers (594 nm, 650 nm), depletion laser (775 nm), acousto-optic modulator (AOM), polarization-maintaining single-mode fibers (curvy orange lines), multi-mode fibers (curvy grey lines), half-wave plate (HWP), quarter-wave plate (QWP), spatial light modulator (SLM), scanning mirror module (including one resonant mirror and two galvanometer mirrors), polarizing beam splitter (PBS), dichroic mirrors, lenses, mirrors, bandpass filters, avalanche photo diodes (APDs), and control electronics. (**b**) CAD rendering of the isoSTED instrument. (**c**) Transmission spectra of the detection bandpass filters and the custom-made quad-bandpass dichroic mirror (highlighted in yellow in (**a**)).

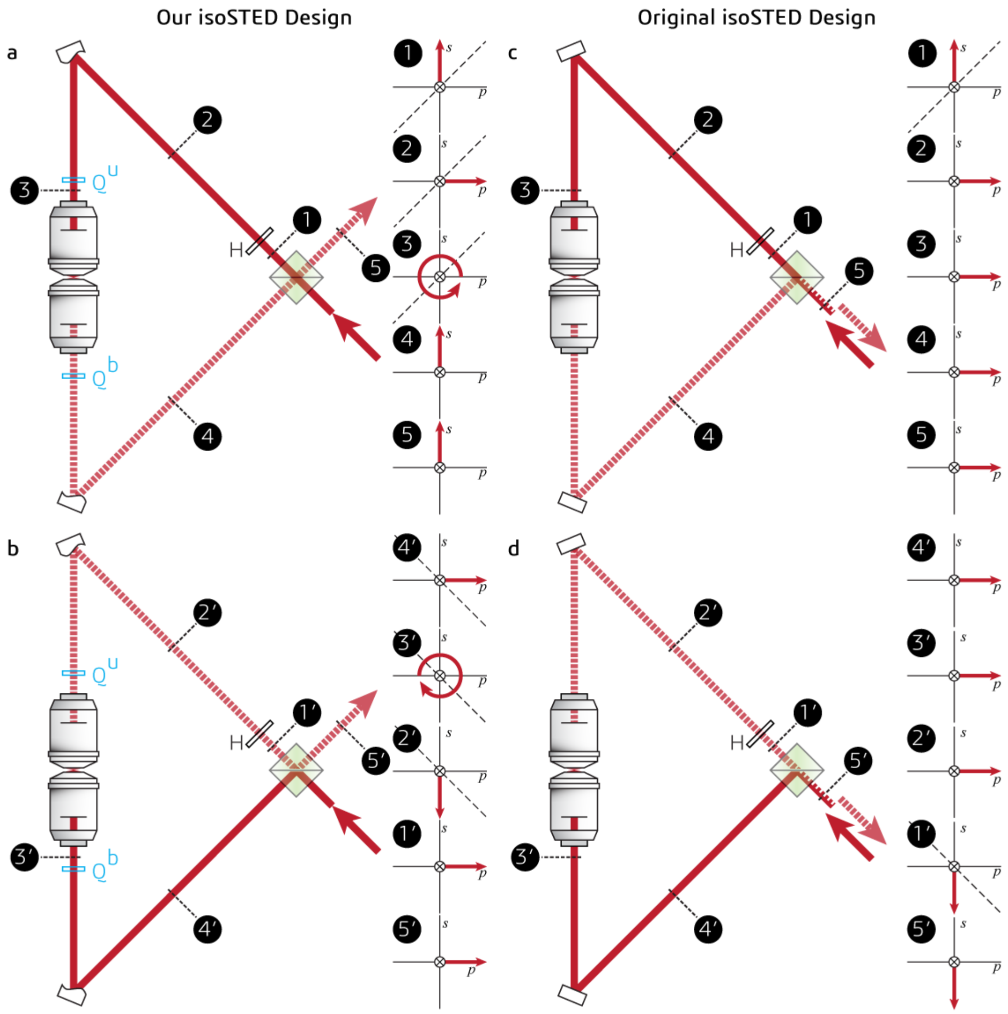


**Supplementary Fig. S2 | Comparison of the polarization states of the depletion beam at indicated points between our setup and the original isoSTED by Schmidt *et al.*** The depletion beam enters the 4Pi interference cavity from the bottom-right side of the PBS. In our design with quarter-wave plates, for the upper arm in the cavity (**a**), H rotates the polarization orientation of the incoming light (solid red line) by 90°. The polarization is further converted to circular by $Q^u$ before entering the upper objective. After propagating through the sample, the outgoing light (dashed red line) is collected by the opposing objective, and its polarization is converted back to linear by $Q^b$. As the polarization orientation is the same as the original one, the beam is transmitted by the PBS and leaves the cavity at the empty (top-right) side of the PBS. Similarly, the incident beam entering the lower arm (**b**) is also dumped to the same side of the PBS. In contrast, in the initial isoSTED nanoscope (**c - d**), the depletion beam leaves the 4Pi cavity at the bottom-right side of the PBS and re-enters the common beam path. On the right of each panel, the polarization at each labelled position is shown. Please note, that these plots are oriented such that the laser beam goes into the image plane from the perspective of the reader. **Abbreviations:** H: half-wave plate; $Q^u$: upper quarter-wave plate; $Q^b$: bottom quarter-wave plate; PBS: polarizing beam splitter.

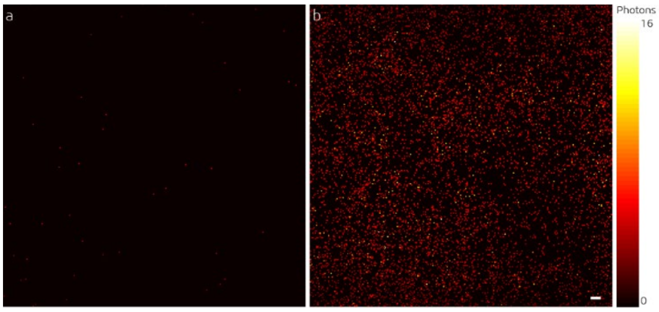


**Supplementary Fig. S3 | Comparison of the background.** The images are respectively taken using the isoSTED nanoscope with (**a**) and without (**b**) quarter-wave plates with the STED laser being on. Scale bar: 1 μm.

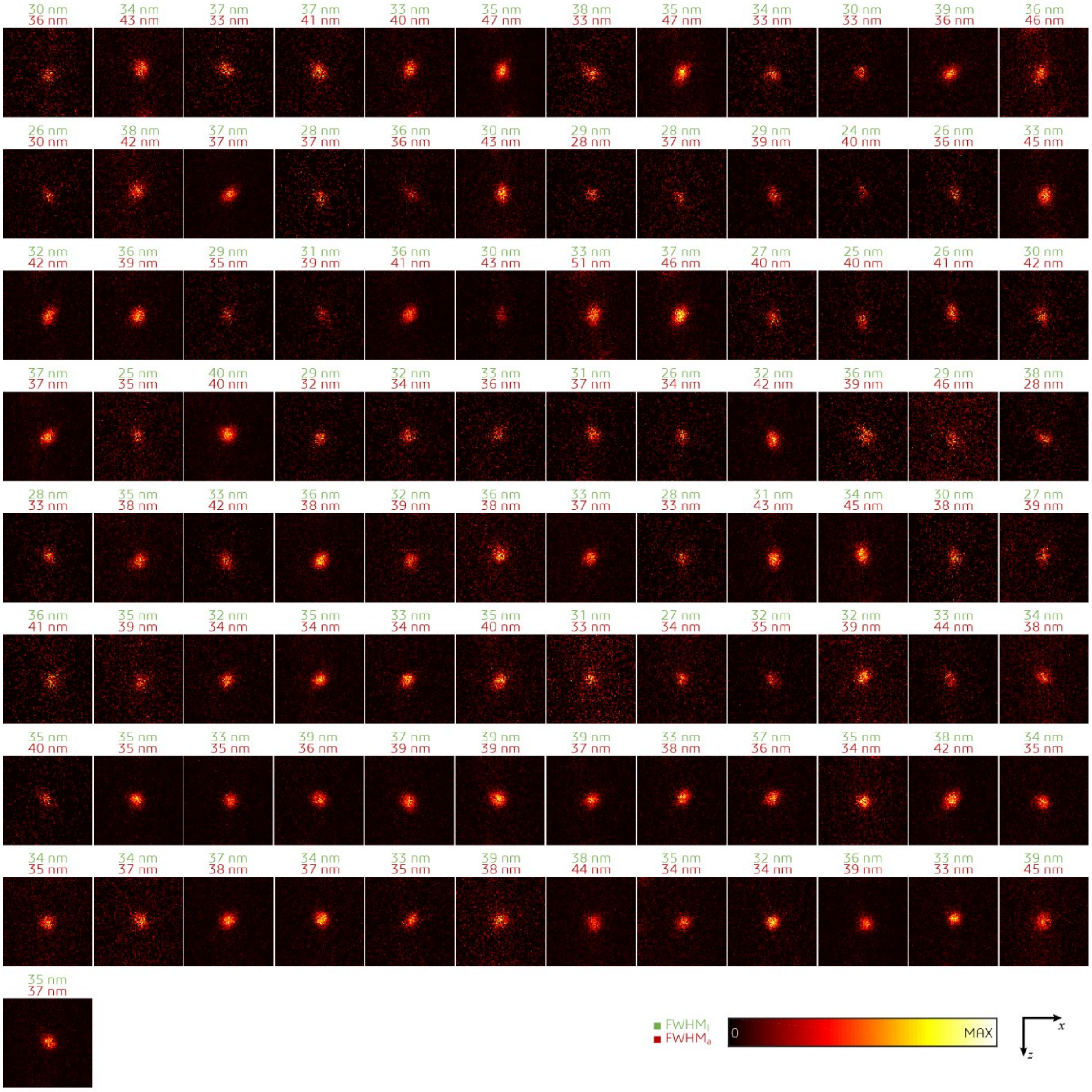


**Supplementary Fig. S4 | Quantification of 3D resolution.** Crimson fluorescent beads with 20 nm diameter were imaged in isoSTED mode. One-dimensional Lorentzian functions were fitted to images of isolated beads in both *x* and *z* directions. The full-width-half-maximum (FWHM) of the fitted functions are marked above each bead image. The numbers in green and red indicate the corresponding FWHMs in lateral (*x*) and axial (*z*) directions, respectively. All images are normalized to their peak intensities.

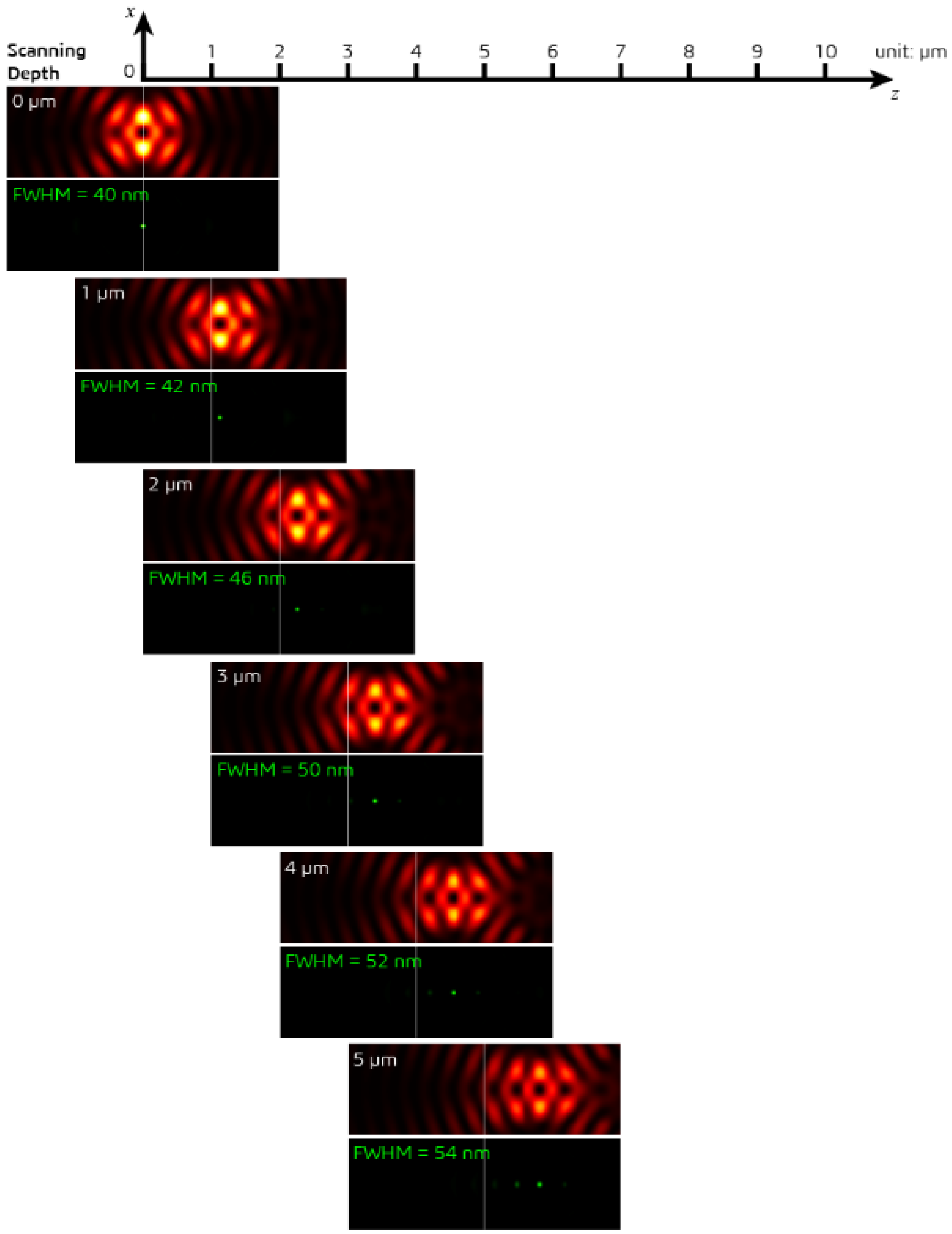


**Supplementary Figure S5 | Effects of refractive index mismatch on the depletion and effective PSF.** The presence of refractive index mismatch between the silicone oil ($n_s$) and the sample mounting medium ($n_i$) causes the appearance of depth dependent side-lobes in the effective PSFs, even when correcting for piston. In addition, the main lobe of the effective PSF also shifts as a function of depth. All results are calculated for $n_s$= 1.406 and $n_i$ = 1.33.

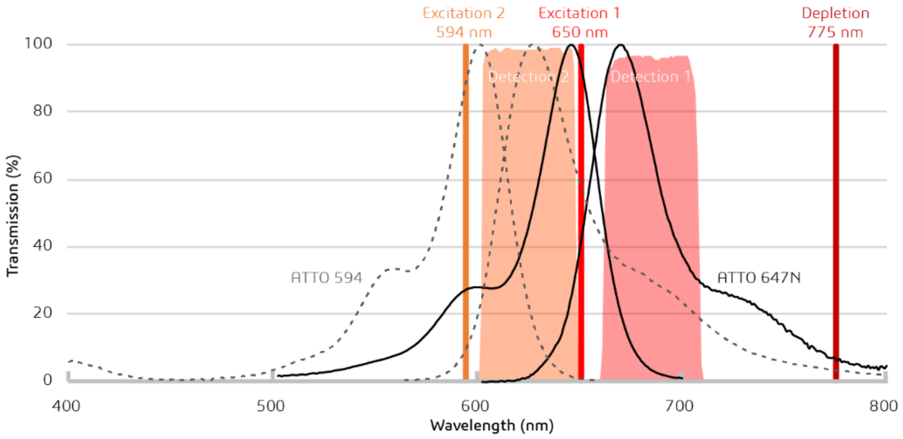


**Supplementary Fig S6 | Diagram of used laser wavelengths, dye emission spectra, and detection filter spectra.** In this diagram, the solid and dashed black curves are the excitation and emission spectra of the dye combination used in our isoSTED nanoscope (solid: ATTO 647N, dashed: ATTO 594). The vertical lines represent the wavelengths of the excitation and the depletion lasers. The transmission of the band-pass filters in front of the detectors (APDs) are shown as the translucent areas.

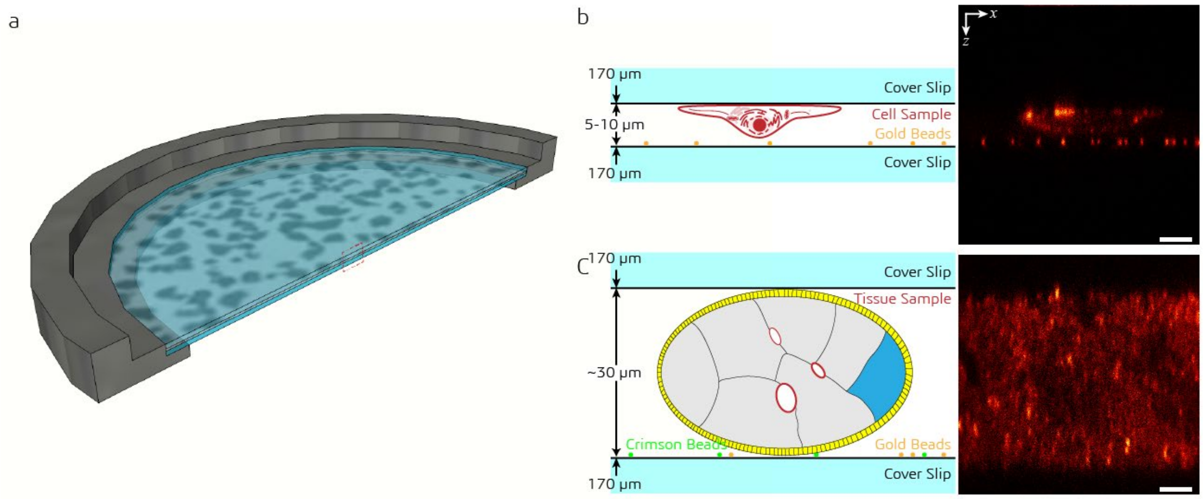


**Supplementary Fig. S7 | Sample holder design.** The specimens are mounted on a ring-shaped sample holder (**a**), between two coverslips at 5-10 µm (**b**, cells) or ~30 µm (**c**, tissue) distance. 100-nm gold beads are sparsely deposited onto the bottom coverslip for use during the aberration correction. Scale bar: 5 µm.

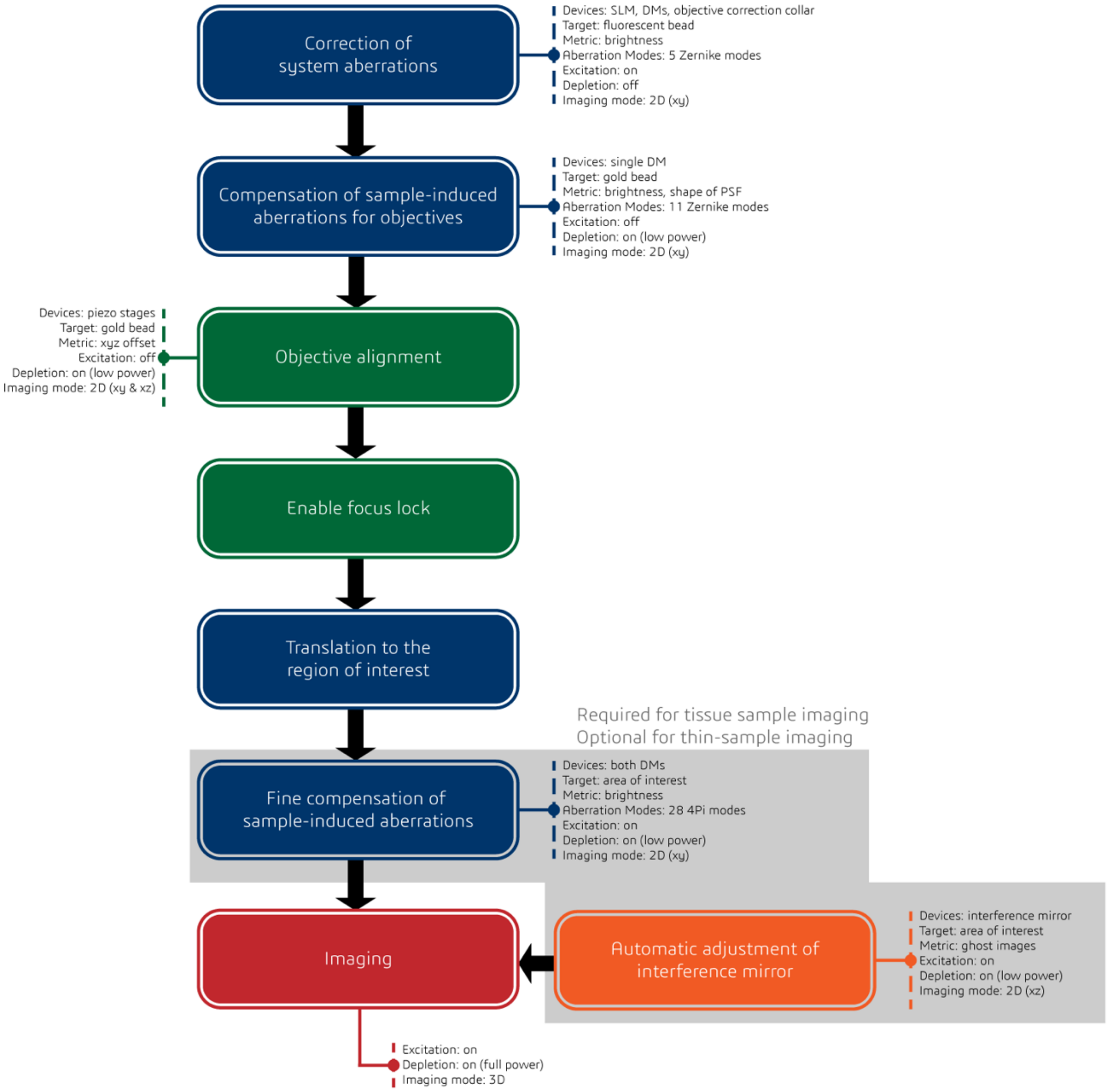


**Supplementary Figure S8 | Detailed process chart of the AO strategy used for tissue imaging via isoSTED nanoscopy.**
